# Highly Sensitive Identification of Lymphatic and Hematogenous Metastasis Routes of Novel Radiolabeled Exosomes Using Non-invasive PET Imaging

**DOI:** 10.1101/2020.03.17.995860

**Authors:** Kyung Oh Jung, Young-Hwa Kim, Seock-Jin Chung, Keon Wook Kang, Siyeon Rhee, Guillem Pratx, June-Key Chung, Hyewon Youn

**Affiliations:** Department of Nuclear Medicine, Medical Research Center, Seoul National University College of Medicine, Seoul, Korea; Biomedical Sciences, Medical Research Center, Seoul National University College of Medicine, Seoul, Korea; Cancer Research Institute, Medical Research Center, Seoul National University College of Medicine, Seoul, Korea; Institute of Radiation Medicine, Medical Research Center, Seoul National University College of Medicine, Seoul, Korea; Tumor Microenvironment Global Core Research Center, Seoul National University, Seoul, Korea; Cancer Imaging Center, Seoul National University Hospital, Seoul, Korea; Department of Nuclear Medicine, National Cancer Center, Goyang, Korea; Department of Radiation Oncology, Division of Medical Physics, Stanford University School of Medicine, Stanford University, Stanford, California, USA; Department of Biology, Stanford University, Stanford, California, USA

**Keywords:** Exosome, Biodistribution, PET image, NIR image

## Abstract

Clinically, there has been significant interest in the use of exosomes for diagnostic applications as promising biomarkers and therapeutic applications as therapeutic vehicles. However, knowledge of *in vivo* physiological biodistribution of exosomes was difficult to assess until now. Physiological distribution of exosomes in the body must be elucidated for clinical application. In this study, we aimed to develop reliable and novel methods to monitor biodistribution of exosomes using *in vivo* PET and optical imaging.

**Methods:** Exosomes were isolated from cultured medium of 4T1, mouse breast cancer cells. Exosomes were labeled with Cy7 and ^64^Cu (or ^68^Ga). In mice, radio/fluorescent dye-labeled exosomes were injected through the lymphatic routes (footpad injection) and hematogenous metastatic routes (tail vein injection). Fluorescence and PET images were obtained and quantified. Radio-activity of *ex vivo* organs was measured by gamma counter.

**Results:** PET signals from exosomes in the lymphatic metastatic route were observed in the draining lymph nodes, which are not distinguishable with optical imaging. Immunohistochemistry revealed greater uptake of exosomes in brachial and axillary lymph nodes than inguinal lymph node. After administration through the hematogenous metastasis pathway, accumulation of exosomes was clearly observed in PET images in the lungs, liver, and spleen, showing results similar to *ex vivo* gamma counter data.

**Conclusion:** Exosomes from tumor cells were successfully labeled with ^64^Cu (or ^68^Ga) and visualized by PET imaging. These results suggest that this cell type-independent, quick, and easy exosome labeling method using PET isotopes could provide valuable information for further application of exosomes in the clinic.

---

**E**xosomes are cell-derived extracellular vesicles containing many functional proteins, mRNAs, and miRNAs (1–3), which are considered as novel messengers in cell-to-cell communication (4,5). Recently, there is growing interest in the clinical application of exosomes for diagnosis, prognosis, and therapy (6–8). Molecular components of exosomes have received increased attention as promising biomarkers for clinical tumor profiling. Exosomes are currently being investigated as therapeutic carriers for drug delivery (9–12). Utilizing exosomes as drug carriers could overcome the limitations of liposomes and nanoparticles, due to their desirable features of exosomes such as highly stable under physiological conditions, non-toxicity, and non-immunogenic. Therefore, the biodistribution of exosomes should be elucidated to further evaluate the efficacy of exosome-based therapeutics.

Previous studies have investigated methods to image exosomes *in vivo* by fluorescence imaging, bioluminescence imaging, radioisotope imaging, MRI imaging, and recent magnetic particle imaging (13–16). Lipophilic fluorescent tracers such as DiR have been used for optical imaging (9,10). Reporter vectors encoding green fluorescent protein conjugated exosome-specific proteins have been used for exosome imaging (17,18). Through these imaging methods, it has been shown that the exosomes localize to the lungs, liver, spleen, and lymph nodes (19). Although these optical exosome labeling methods are useful to visualize the localization of exosomes *in vitro* and *ex vivo* studies (20–22), it is still difficult to quantify the biodistribution of exosomes *in vivo* due to low tissue penetration and low sensitivity.

On the other hand, radionuclide imaging has been suggested as an option to quantify imaging signals *in vivo* and overcome penetration depth limitations with better sensitivity (23). Recently, radioisotopes such as ^111^In, ^125^I, and ^99m^Tc-HMPAO were used to monitor the biodistribution of exosomes (24–26). However, ^111^In or ^125^I-labeled exosome imaging could only provide insight into *ex-vivo* biodistribution using gamma counter. Moreover, the deiodination of the radioiodine labeled ligand in the case of ^125^I labeling has been reported as a drawback to its *in vivo* application (27). Although ^99m^Tc-HMPAO-based exosome imaging was used to visualize *in vivo* exosome biodistribution, it is also limited by the glutathione (GSH)-dependent accumulation of ^99m^Tc-HMPAO in exosomes. The level of GSH in exosomes varies between cell types and can be easily altered by various enzymes in the cells (28). Furthermore, ^99m^Tc-HMPAO imaging can only be applied to single-photon emission computed tomography (SPECT), which is not as sensitive and quantitative as positron emission tomography (PET). In addition, the standards for PET image acquisition and the methods for quantitative data analysis of PET images have already been established by the nuclear imaging society (29).

In this study, we developed a simple and easy exosome radiolabeling method that is less cell-type dependent and utilize more quantitative PET imaging to better visualize the biodistribution of exosomes in mice. Since 1,4,7-triazacyclononane-triacetic acid (NOTA) is a useful chelator for various radioisotopes, we used NOTA to conjugate with the amine group of the membrane proteins on exosomes (Fig. 1). Cancer cells can metastasize through well-known processes, including lymphatic or hematogenous spread. Exosomes derived from cancer cells could be transferred through the same routes, where they become involved in tumor growth, immune suppression, and metastasis (30,31). Therefore, we investigated the biodistribution of exosomes through these routes with quantitative PET imaging which provides valuable information for their clinical application (32,33).

**FIGURE 1.**
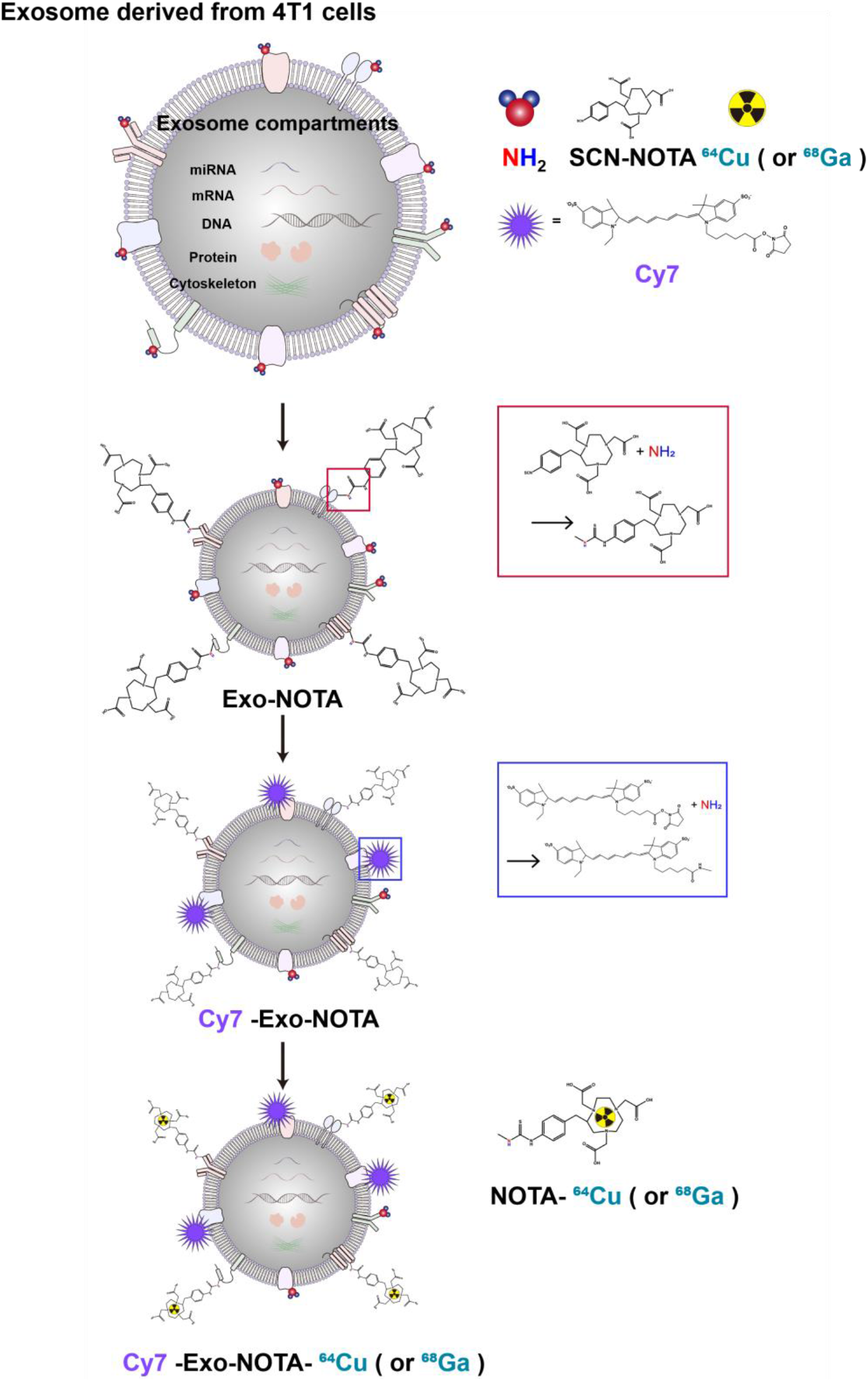
Experimental scheme of exosome labeling. Sequential exosome labeling steps of SCN-NOTA as a chelator for radio-isotope, Cy7 fluorescence dye, ^64^Cu and ^68^Ga for PET and optical imaging.

## MATERIALS AND METHODS

### Cell Lines

The 4T1 mouse mammary tumor cells were cultured in RPMI 1640 medium (Invitrogen, Grand Island, NY, USA) supplemented with 10% fetal bovine serum and 1% penicillin and streptomycin mix (Invitrogen, Grand Island, NY, USA).

### Exosome Purification

Exosomes were purified from cultured medium supplemented with exosome-depleted FBS using the exosome purification kit following manufacturer’s instructions (ExoQuick™, System Bioscience, Mountain View, CA, USA).

### Western Blotting

Extracted proteins (20 μg) of exosomes incubated with antibodies for anti-CD9, anti-CD63 (System Bioscience, Mountain View, CA, USA), anti-AIP1/Alix (BD Biosciences, San Jose, CA, USA), and anti-calnexin (Santa Crux Biotechnology, Santa Cruz, CA, USA). The following secondary antibodies were incubated. Immunoreactive bands were imaged with a LAS-3000 imaging system (Fuji Film, Tokyo, Japan).

### Transmission Electron Microscopy (TEM)

The exosomes were fixed with 2% glutaraldehyde at 4°C and deposited using a copper grid (300 mesh and covered with carbon). The morphology and size of the exosomes was imaged with a JEM 1400 transmission electron microscope (JEOL, CA, USA).

### Synthesis of Exosome-NOTA

One hundred microliters of sodium carbonate buffer (1 M, pH 9.5) was added to 100 μg of exosomes and mixed with 1 μL of *p-*SCN-Bn-NOTA (200 μg/μL, Macrocyclics, Inc., TX, USA). The reaction mixture was incubated for 1 h at 4°C. Twenty microliters of ExoQuick™ was added to purify exosome-NOTA.

### Fluorescent Labeling of Exosome

The exosomes (100 μg) were incubated with Cy7 mono-NHS ester (5 μM, Sigma-Aldrich, MO, USA), a near-infrared fluorescence dye, for 10 min at 37°C (23). Cy7-labeled exosomes were purified using ExoQuick™.

### ^64^Cu Labeling of Exosome

Five hundred microliters of sodium acetate buffer (2 M, pH 5.2) was added to a 1.5 mL tube containing 100 μg of exosome. Subsequently, exosomes were labeled with ^64^Cu (7.4 MBq) by adding 100 μL of a CuCl_2_ solution in 0.1 N HCl (KIRAMS, Seoul, Korea). The reaction mixture was incubated under gentle shaking conditions at 37°C for 5 min. The radioactivity was analyzed by using instant thin-layer chromatography silica gel with chromatography paper as a stationary phase and citrate acid buffer (0.1 M, pH 5.0) as a mobile conjugate. To remove unconjugated free-^64^Cu, ExoQuick™ was added to ^64^Cu labeled exosomes for purification.

### *In Vitro* Serum Stability Test

The stability of exosome-NOTA-^64^Cu in the serum was tested by incubating them in 10% of exosome-depleted FBS at 37°C until 36 h. Radiolabeling efficiency was measured by ITLC-SG analysis.

### ^68^Ga Labeling of Exosome

One hundred microliters of sodium acetate (2 M, pH 5.2) was added to a 1.5 mL tube containing 100 μg of exosome. Subsequently, exosomes were labeled with ^68^Ga (33.3 MBq) by adding 400 μL of a GaCl_3_ solution. The reaction mixture was incubated under gentle shaking conditions at 25°C for 30 min. Radioactivity was determined using ITLC-SG. ExoQuick™ was added to purified the exosomes.

### *In Vivo* Mouse Study

All procedures for the *in vivo* studies were approved by the Institutional Animal Care and Use Committee in Seoul National University Hospital (SNUH-IACUC) and adhered to the Guide for the Care and Use of Laboratory Animals. The 8-week-old female BALB/c nu/nu mice were used for *in vivo* studies weighing about 20 g on average.

### *In Vivo* Exosome Injection in Mice

Cy7- and ^64^Cu- or ^68^Ga-labeled exosomes (20 μg) were intravenously injected into the tail vein or subcutaneously injected through the footpad. Fluorescence and PET images were acquired.

### *In Vivo* and *Ex Vivo* Fluorescence Imaging

The Cy7 signals from exosomes were imaged using the IVIS100 imaging system (Xenogen Corp., Alameda, CA, USA). For tissue imaging, the Zeiss LSM510 META confocal imaging system (Carl Zeiss, Thornwood, CA, USA) was used.

### *In Vivo* PET Imaging

The ^64^Cu signals from exosomes were imaged using the Genesys4 (Sofie Bioscience, Inc., CA, USA). The radioactivity of the exosomes was around 0.74 MBq at the time of the injection. The PET images were acquired for 5 min with the x-ray setting set at 100 μA, 40 kVp, and with a 3 sec exposure. Image data was automatically reconstructed by using a 3D MLEM algorithm, which was evaluated by A region of interest (ROI) analysis with the AMIDE software package. ROIs were drawn over the target organ margin with results expressed standardized uptake value (SUV).

### Radioactivity for *Ex Vivo* Tissue Samples

Radioactivity associated with each organ was measured with a gamma counter (Packard, Meriden, CT, USA) and was expressed as percentage of injected dose per gram of tissue (%ID/g) for a group of 5 animals.

### Immunohistochemistry

Tissues were stained with antibody for anti-CD63 (System Bioscience, Mountain View, CA, USA). The secondary antibody, biotinylated anti-rabbit for CD63 (Dako, Glostrup, Denmark), was incubated. To confirm the uptake of exosomes in immune cells, the anti-CD63 fluorescent antibody was co-stained with CD11b (Abcam, Cambridge, MA, UK), F4/80 antibody (Abcam, Cambridge, MA, UK), and CD90.2 antibodies (BD Biosciences, San Jose, CA, USA).

### Statistical Analysis

At least three independent samples were tested in each group, and data were expressed as mean ± SD and statistical significance determined using the One-way ANOVA (and nonparametric). The ANOVA was used for multiple comparisons (GraphPad Soft ware Inc., La Jolla, CA, USA). P < 0.05 was considered as statistically significant.

## RESULTS

### Exosome Characterization and Labeling with Cy7, ^68^Ga, and ^64^Cu

In purified exosomes, exosome marker proteins such as CD9, CD63, and Alix were expressed higher in exosomes than in cells, while cell marker proteins, such as calnexin, were expressed more highly in cells (Fig. 2A). We confirmed the presence of extracelluar vesicles of the appropriate size (about 100 nm) using DLS and morphology of exosomes through TEM (Fig. 2B, C). Shown in Fig.1, our labeling strategy is based on the use of SCN-NOTA as a bifunctional chelator of radioisotopes such as ^64^Cu and ^68^Ga. We also labeled with Cy7, a near infra-red fluorescence dye, for optical imaging. Cy7-labeled exosome showed clear fluorescence signals in the tube (Fig. 2E). Thin layer chromatography, commonly used to confirm serum stability of radiolabeled tracer, demonstrated that after labeling with radioisotopes, the ^64^Cu/^68^Ga-labeled exosomes (Exo-NOTA-^64^Cu/^68^Ga) had a labeling purity of approximately 98% after removal of free radioisotope by ExoQuick (Supplemental Fig. 1). The stability of radiolabeled-exosomes was tested in the serum. ^64^Cu-labeled exosomes (Exo-NOTA-^64^Cu) in serum were stable until 36 h after labeling (Fig. 2F).

**FIGURE 2.**
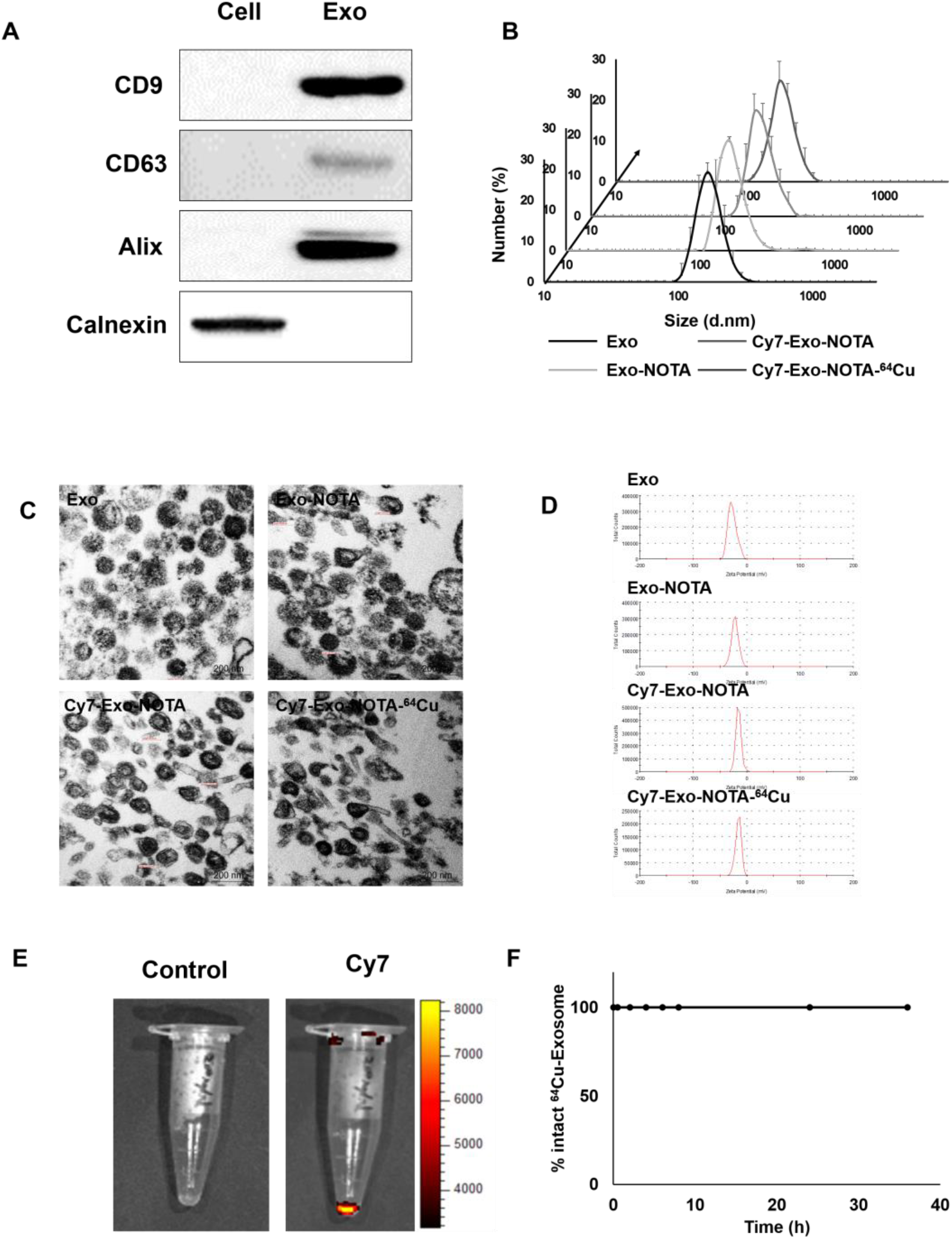
Characterization and labeling of exosomes. (A) Western blot analysis of common exosome markers (CD9, CD63 and Alix) and cell marker (Calnexin). (B) The size distribution of 4T1 derived exosome population according to labeling steps. (C, D) Representative transmission electron microscopy (TEM) images and zeta-potential of exosomes according to labeling steps. (E) Fluorescence Imaging of Cy7-labeled Exosomes compare to the control exosomes. (F) Serum stability test of exosome-^64^Cu until 36 h.

### *In Vivo* Exosome Uptake Imaging in the Lymphatic Route

PET images at 24 h after lymphatic injection of exosomes showed that radiolabeled exosomes (Exo-NOTA-^64^Cu) had greater uptake in lymph nodes than NOTA-^64^Cu (Fig. 3A) and Free-^64^Cu (Supplemental Fig. 2A). In whole body PET images, there is no significant uptake in other organs (Supplemental Fig. 2D, E). Optical images showed that Cy7 signals from exosomes were detected only in the brachial lymph node and at the injection site (Fig. 3A), whereas PET images could clearly visualize the localization of exosomes in the axillary lymph node as well as in the brachial lymph node with higher sensitivity. For ^68^Ga-labeled exosomes, the results were similar to those for exosome-^64^Cu. ^68^Ga-labeled exosomes (Exo-NOTA-^68^Ga) showed greater accumulation in the lymph nodes than NOTA-^68^Ga at 1 h after injection (Supplemental Fig. 3).

**FIGURE 3.**
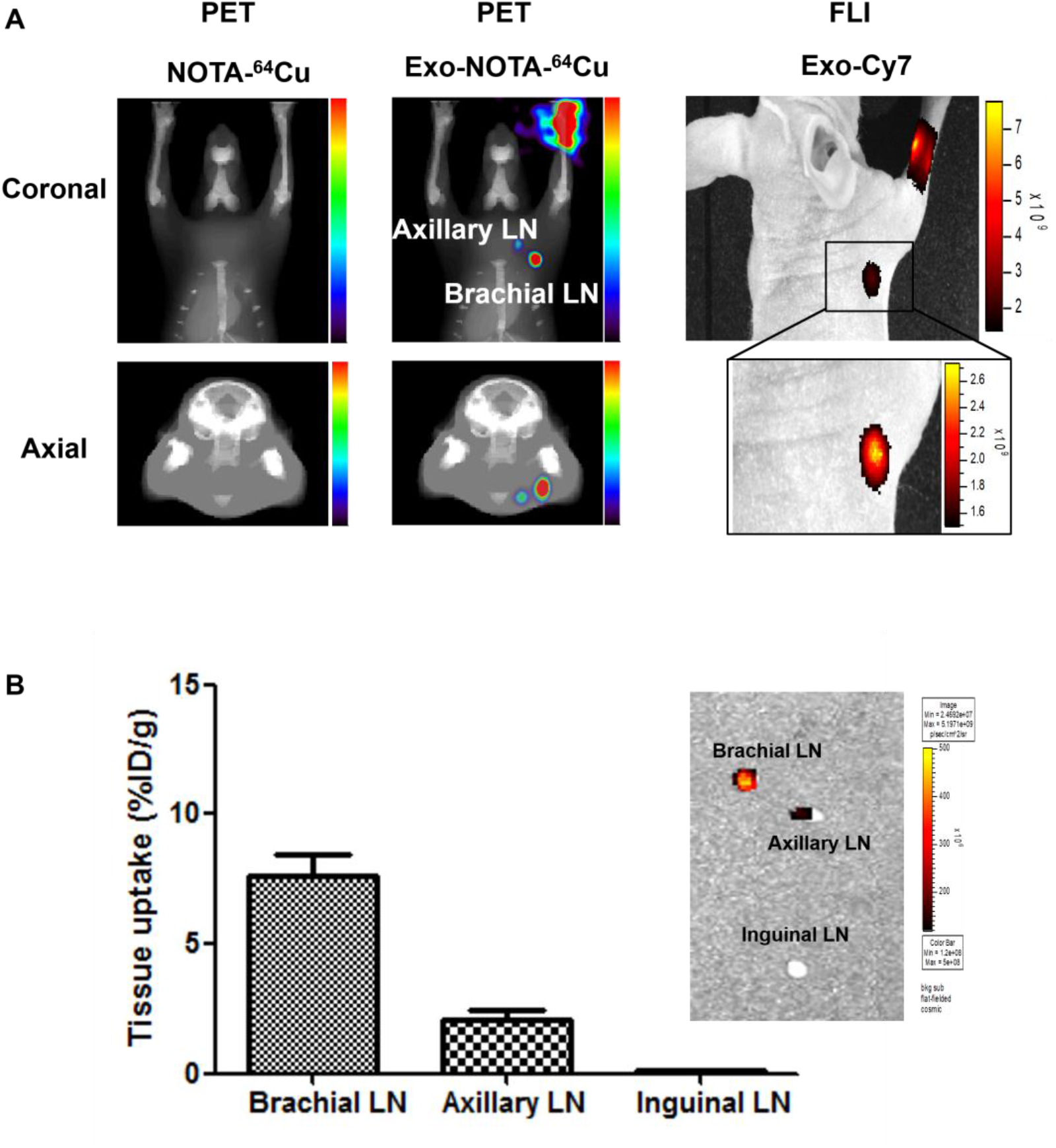
*In vivo* imaging of exosomes in the lymphatic route. (A) Footpad injected exosomes (Exo-NOTA-^64^Cu) have more uptake in the lymph nodes than NOTA-^64^Cu. ^64^Cu signals of exosomes are detected in the brachial and axillary lymph nodes with higher sensitivity than fluorescence imaging. (B) *Ex vivo* fluorescence images show Cy7 signals of exosomes in the brachial and axillary lymph nodes. Radioactivity in brachial and axillary lymph nodes is stronger than that in the inguinal lymph nodes. Data represent mean ± s.d. (n = 5/group).

### *Ex Vivo* Exosome Uptake Imaging in the Lymphatic Route

In surgically removed lymph nodes, Cy7 signals were high in the brachial and axillary lymph nodes, similar to what was observed on the PET images (Fig. 3B). The radioactivity in gamma counter was stronger in the brachial and axillary lymph nodes than in the inguinal lymph node. The expression of CD63, an exosome marker, was higher in the exosome-injected mice than non-injected mice (Fig. 4). CD63 expression was higher in the brachial and axillary lymph nodes than in the inguinal lymph node. Confocal microscopy images also showed that stronger Cy7 signals were observed in the brachial and axillary lymph nodes than in the inguinal lymph node.

**FIGURE 4.**
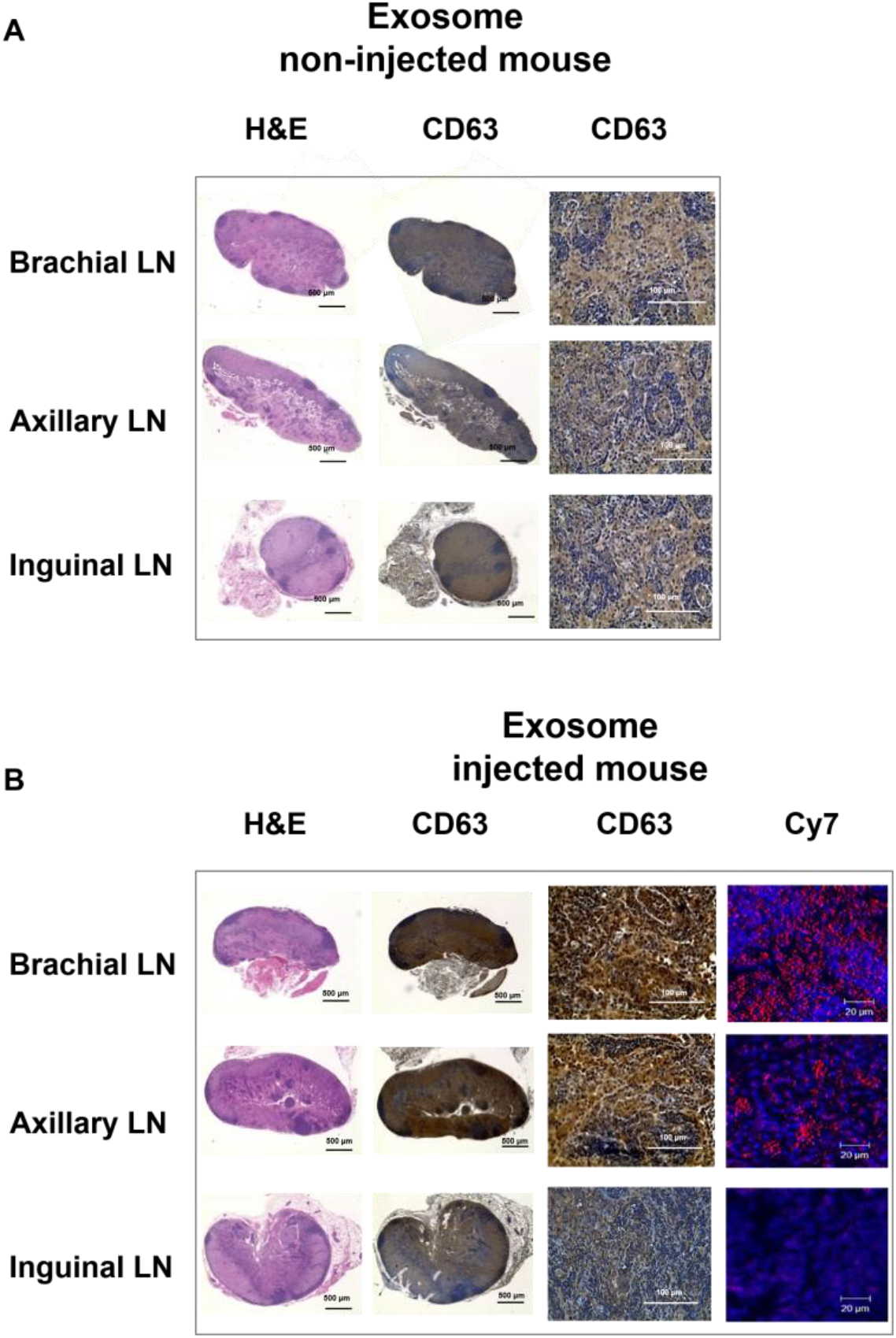
*Ex vivo* imaging of exosomes in the lymphatic route. The expression of CD63 (as an exosome marker) in lymph node is higher in mice injected with exosomes (A) than in mice that are not injected mice (B), with higher brachial and axillary lymph nodes, compared to the inguinal lymph nodes. A confocal microscope images show stronger Cy7 signals in the brachial and axillary lymph nodes.

### *In Vivo* and *Ex Vivo* Imaging in the Hematogenous Route

PET image revealed that ^64^Cu-labeled exosomes (Exo-NOTA-^64^Cu) had accumulated in the lungs and liver at a greater rate than NOTA-^64^Cu (Fig. 5A) and Free-^64^Cu (Supplemental Fig. 4). Intravenously injected exosomes mainly accumulated in the lungs and liver at the initial time point, after which they circulated and were cleared from the blood. However, Cy7 fluorescence signals in mouse were not visible (Fig 5B, left), whereas the Cy7 signals from *ex vivo* organs showed strong signals from lungs and liver similar to PET images (Fig 5B, right), highlighting the greater sensitivity of PET imaging. For evaluating the *in vivo* biodistribution of systemically injected exosomes in PET images, a ROI was automatically drawn over the target organ margin based on the CT image. In addition, radioactivity of exosomes from *ex vivo* organs were also quantified by gamma counter. Biodistribution of exosomes determined by quantifying PET imaging data (Fig. 6A) was almost the same as those determined from the *ex vivo* radioactivity (Fig. 6B), indicating that PET imaging with ^64^Cu-labeled exosome provide quantitative information. For ^68^Ga-labeled exosomes, the results showed the same pattern as that of ^64^Cu-labeled exosomes, showing strong signals from lungs and liver (Supplemental Fig. 5). The expression of CD63 in the lungs, liver, and spleen was higher in the exosome-injected mice than non-injected mice, not completely same pattern with PET imaging data (Fig. 6C).

**FIGURE 5.**
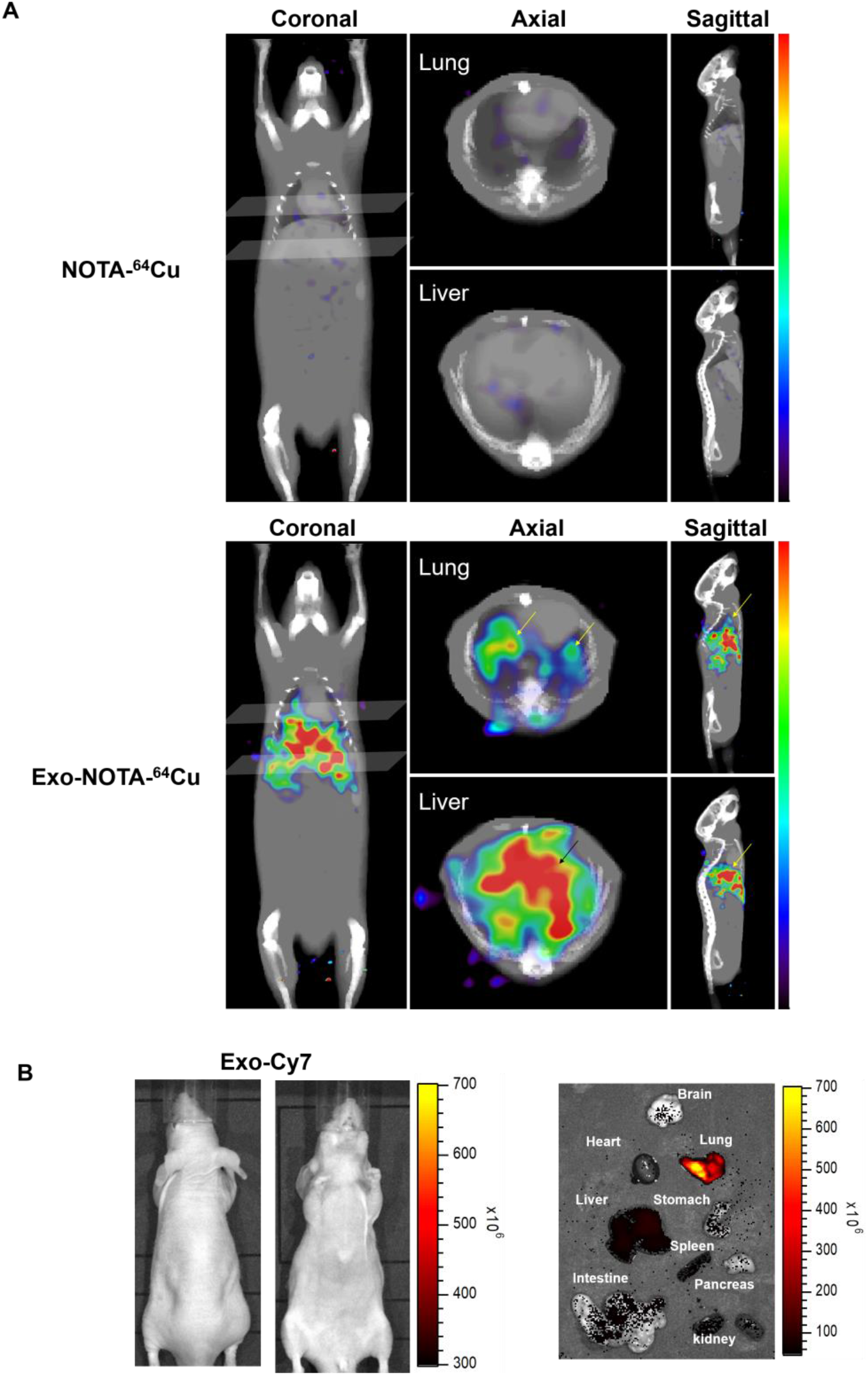
*In vivo* imaging of exosomes in the hematogenous route. (A) Exo-NOTA-^64^Cu was observed to have more uptake in the lungs and liver at 24 h than NOTA-^64^Cu. (B) Exo-Cy7 was not detected in systemic fluorescence images. Only *ex vivo* fluorescence images was shown strong uptake of exosomes in the lungs, liver, and spleen. (n = 5/group).

**FIGURE 6.**
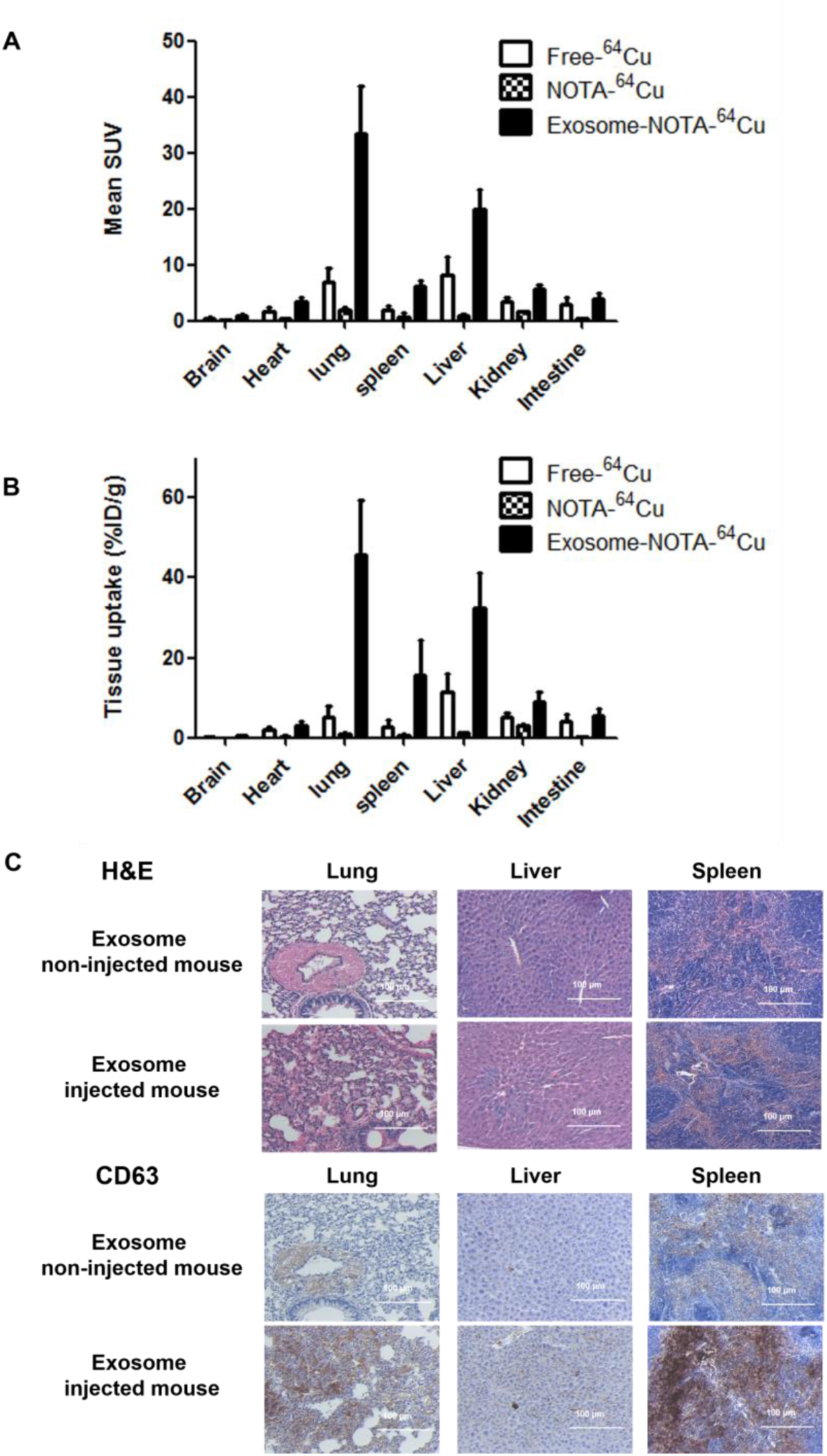
*Ex vivo* imaging of exosomes in the hematogenous route. (A) Biodistribution of exosomes was determined by quantifying PET imaging data. Mean SUV indicates the average of standard uptake value (n = 5). (B) *Ex vivo* organ uptake signals using a gamma counter is similar to signals of PET images. (C) The expression of CD63 is higher in the mice injected with exosomes. Data represent mean ± s.d. (n = 5/group).

## DISCUSSION

In contrast to previous exosome imaging systems (24–26), ^64^Cu− or ^68^Ga-labeling with NOTA-conjugated exosomes is very simple and easy, providing more quantitative information in living animals using PET imaging. This NOTA-conjugated labeling method for exosome amine groups has an advantage in cell type-independent radiolabeling, because most cancer cell-derived exosomes have sufficient amine groups on their surface (1,2).

Furthermore, *in vivo* quantification of PET images from specific organs provided accurate information about the biodistribution of exosomes compared with radioactivity of the *ex vivo* organs. This result indicates that PET images indeed can provide quantitative information about *in vivo* biodistribution, and our exosome labeling method with ^64^Cu or ^68^Ga is an excellent candidate for clinical application. Our exosome labeled with ^68^Ga showed similar distribution with ^64^Cu. Even though ^68^Ga (68 min) has short half-life and low labeling efficiency compared with ^64^Cu (12.7 hr), ^68^Ga is easy-to-use and low-cost for users without cyclotrons. Therefore, ^68^Ga data was included as supplementary information. In addition, various radioisotopes are applicable with NOTA in this system. Because therapeutic radioisotopes such as ^177^Lu (6.7-day half-life) were labeled with NOTA (34,35), it may be possible to monitor exosomes in long-term images and to use exosomes as therapeutic agents. Other chelators such as DOTA and DTPA could be used to label exosomes for MRI as well as another radionuclide imaging (23,36). Therefore, our method could be used for various clinical applications to monitor the physical location of exosomes in the body, especially for the use of exosomes as therapeutic vehicles (12,37,38).

In this study, we visualized the biodistribution of exosomes in lymphatic or hematogenous metastasis routes. Previous publications on the biodistribution of melanoma-derived exosomes showed that exosomes injected through the lymphatic system were localized in the sentinel lymph node, which was confirmed with *ex vivo* fluorescence images (17,19). Even though these fluorescent images were *ex vivo* images, fluorescent imaging has shown great potential (39). However, since it is difficult to acquire *real-time* fluorescence images, our radiolabeled exosome imaging could be a superior method for monitoring exosomes *in vivo,* overcoming the drawbacks of fluorescence imaging such as limited depth penetration. The homing of exosomes in the sentinel lymph nodes can promote formation of a pro-tumorigenic niche for metastasis, changing the extracellular matrix and vascular proliferation (40,41). After that, cancer cells could be recruited to generate secondary metastatic tumors in these lymph nodes (42,43). Other studies showed that exosomes from tumor cells could circulate through the blood and accumulate in specific organs (17,44). However, because that previous study was unable to confirm uptake of exosomes in *real-time* mice, we aimed to confirm the uptake of exosomes *in vivo*. Our results showed that Cy7 signals of exosomes were detected only in the brachial lymph node and the injection site, whereas ^64^Cu signals of exosomes were detected in all draining lymph nodes, such as the brachial and axillary lymph nodes, in addition to the injection site (Fig. 3A). Both fluorescence images and PET images showed exosomes in the lymph nodes, but PET imaging was more sensitive and had better resolution than fluorescence imaging.

PET images also showed that intravenously injected exosomes accumulated in the lungs and liver, which was not detected in optical images, highlighting the greater sensitivity of PET image over optical imaging (Fig. 4A). Due to the limited penetration depth of fluorescence light, the fluorescence images had limited use for *in vivo* applications (23). Additionally, PET images for ^64^Cu-labeled exosomes showed that the exosomes initially accumulated in the lungs and liver. In the case of ^68^Ga-labeled exosomes (Supplementary Fig. 4B), the accumulation of exosomes was similar with exosome-^64^Cu, showing uptake in the lungs and liver. These results indicate that exosomes can accumulate in the lymph nodes, lungs, and liver through lymphatic or hematogenous routes, which is consistent with the known common metastatic sites of breast cancer in previous distribution studies (18,24,45). Since breast cancer primarily metastasizes to the lungs, regional lymph nodes, liver, and bone (46), our exosome accumulation results could indirectly show preferential exosome accumulation at metastatic sites, forming a pre-metastatic niche.

For future studies, our novel imaging systems could be useful for predicting the *in vivo* biodistribution of exosomes released from different types of cells such as cancer cells, immune cells, or stem cells (11,47). For therapeutic application, quantitative PET imaging of exosomes provides valuable information for drug delivery and bioengineering exosomes for tumor targeting could be used to enhance therapeutic effects (48,49). Furthermore, our imaging systems could be helpful to investigate the role of exosomes as cancer vaccine in cancer immunotherapy (50,51)

## CONCLUSION

Cancer cell-derived exosomes play an important role in intercellular communication by mediating the transfer of biological-information. The *in vivo* distribution is essential for clinical use of therapeutic exosomes. In this study, we established a NOTA-based ^64^Cu− or ^68^Ga− labeling method for monitoring the biodistribution of exosomes with PET imaging. Our data successfully demonstrates that footpad-injected exosomes accumulate in the draining lymph nodes, and intravenously injected exosomes accumulate in the lungs, liver, and spleen. This is the first PET imaging of radiolabeled exosomes *in vivo*. Our ^64^Cu (or ^68^Ga) labeling of exosomes is very simple and less cell-type dependent, providing more quantitative information in living animals. Moreover, our strategy may also be applicable to human trials because PET is already available in the clinic. Therefore, this novel imaging system can be useful for predicting the biodistribution of exosomes *in vivo* and tracking therapeutic exosomes for clinical application.

## SUPPLEMENTAL TABLE

**Table 1.** Hydrodynamic size, PDI (poly dispersion index) and zeta potential of exosome according to the labeling steps.

|  | **Exo** | **Exo-NOTA** | **Cy7-Exo-NOTA** | **Cy7-Exo-NOTA-<sup>64</sup>Cu** |
| --- | --- | --- | --- | --- |
| Size (nm) | 160.3 ± 28.06 | 163.7 ± 19.44 | 189.9 ± 21.33 | 185.6 ± 23.18 |
| PDI | 0.306 | 0.370 | 0.494 | 0.457 |
| Zeta (mV) | -21.8 ± 2.95 | -18.8 ± 2.655 | -15.5 ± 1.97 | -14.9 ± 2.79 |

## SUPPLEMENTAL FIGURE

**SUPPLEMENTAL FIGURE 1.**
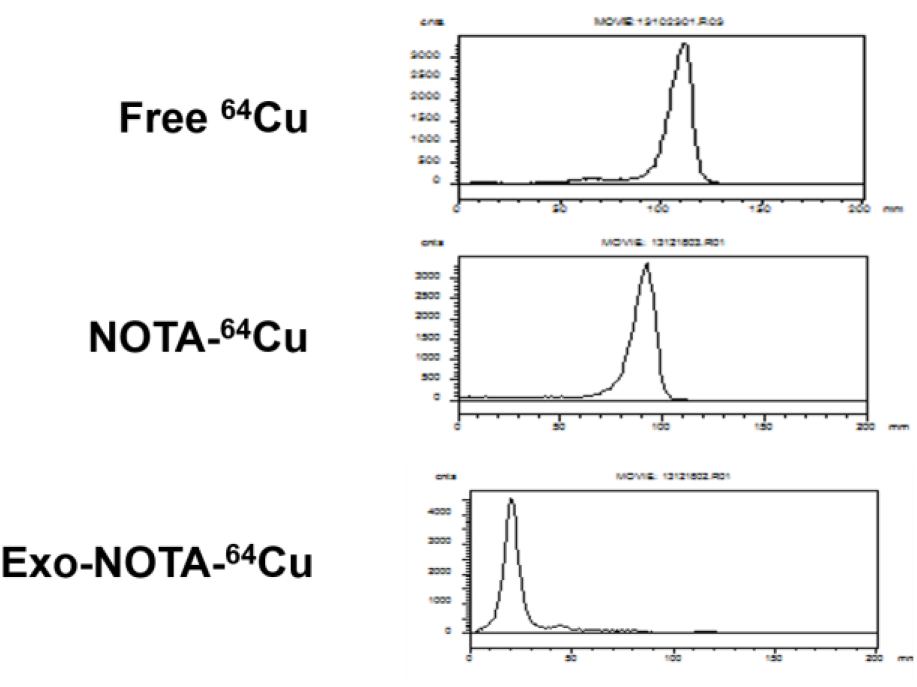
Thin layer chromatography of ^64^Cu. After labeling with radioisotopes, thin layer chromatography shows a labeling efficiency.

**SUPPLEMENTAL FIGURE 2.**
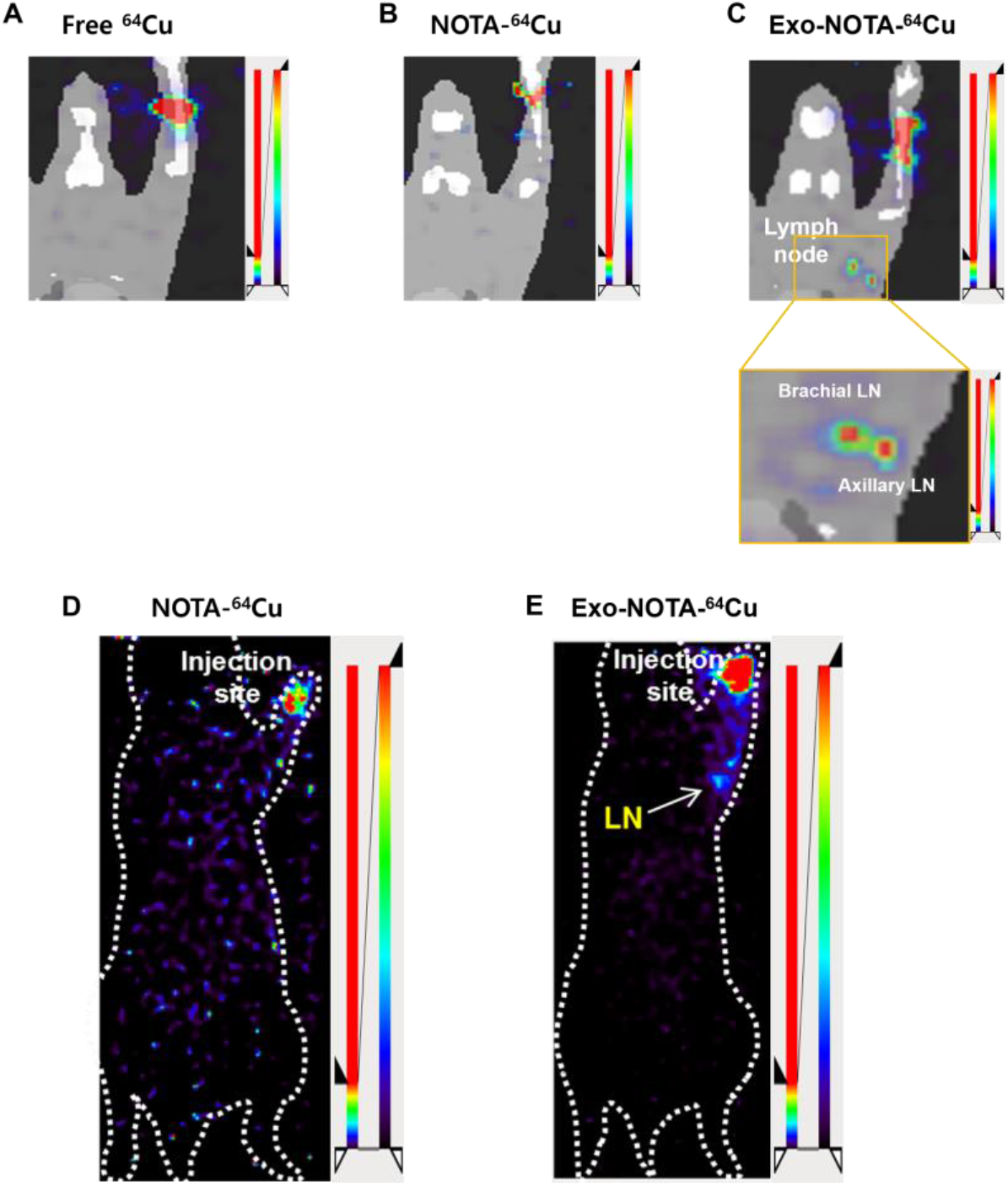
^64^Cu images in the lymphatic route. (A) Free ^64^Cu is compared with NOTA-^64^Cu (B) and Exo-NOTA-^64^Cu (C). (D, E) Whole body PET images showed there is no significant uptake in other organs.

**SUPPLEMENTAL FIGURE 3.**
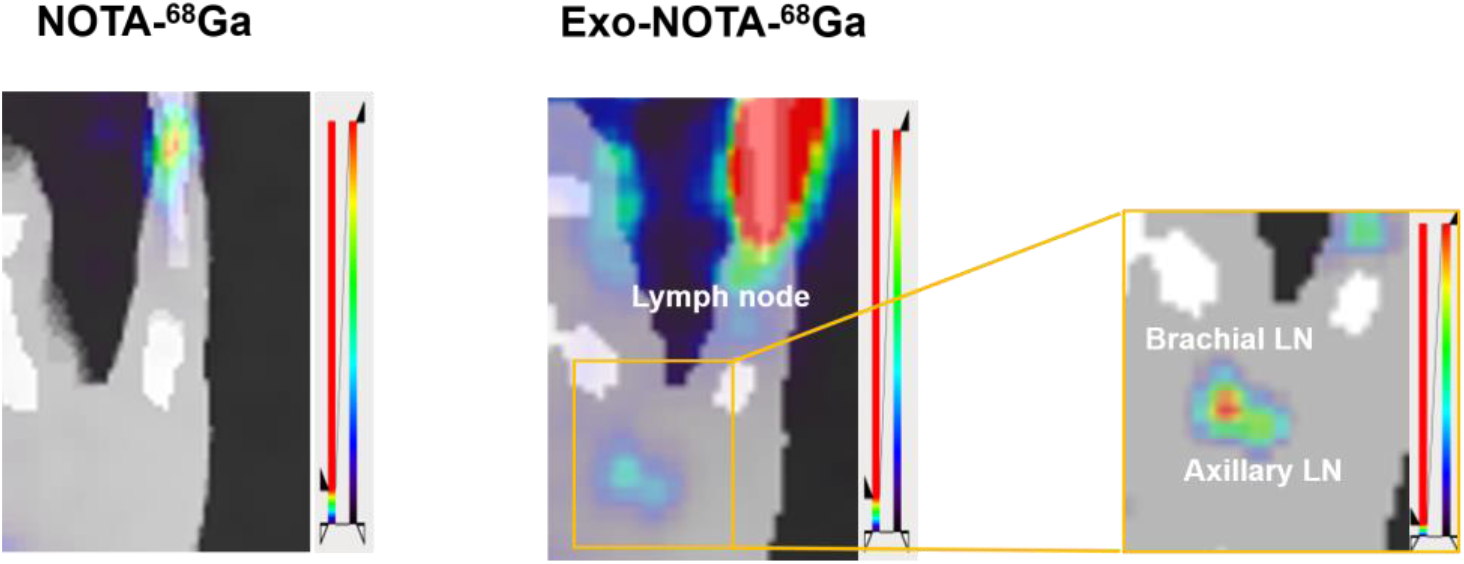
^68^Ga-labeled exosomes in the lymphatic route. Exo-NOTA-^68^Ga is compared with NOTA-^68^Ga.

**SUPPLEMENTAL FIGURE 4.**
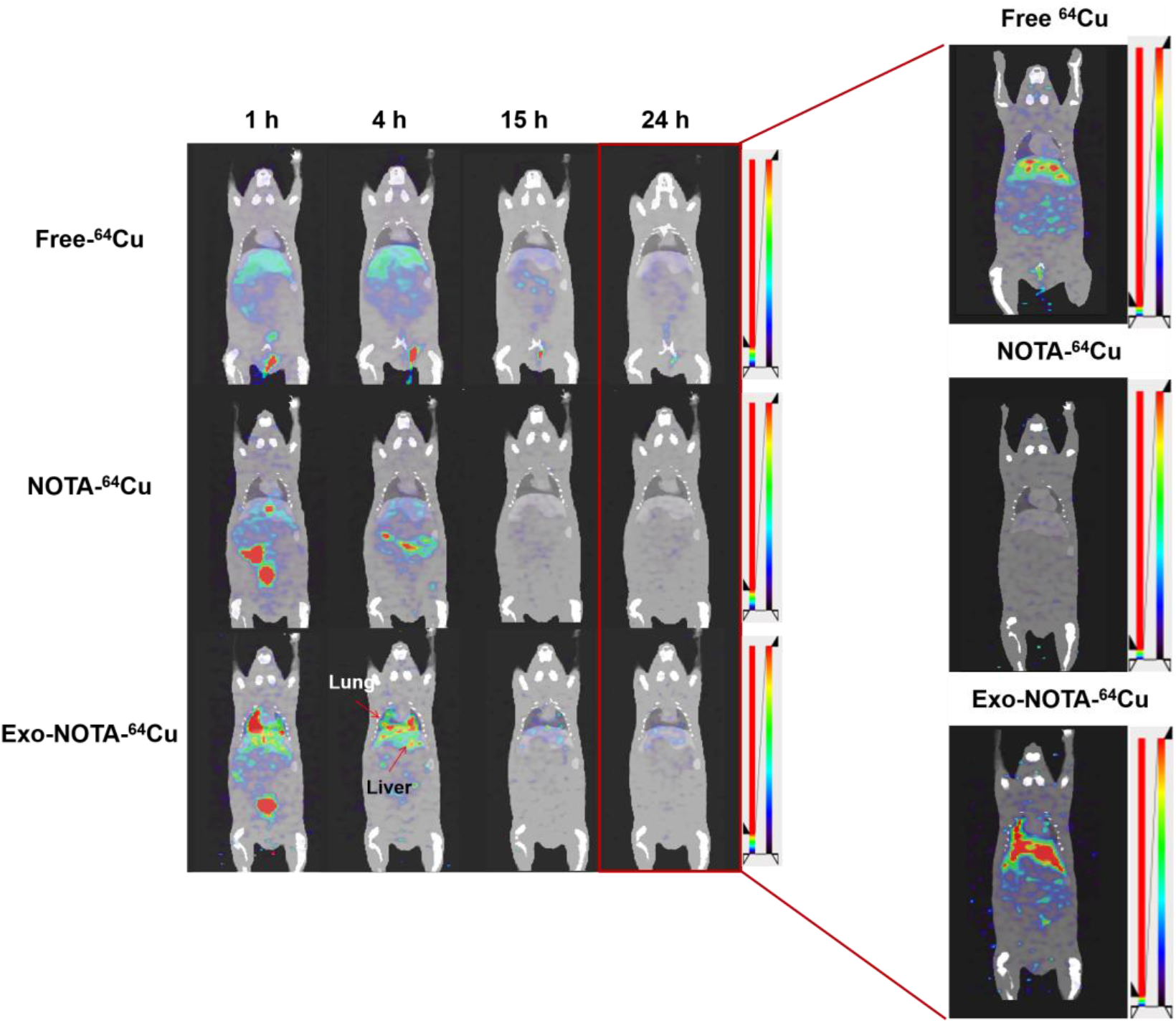
Free-^64^Cu images in the hematogenous route. Free ^64^Cu is compared with NOTA-^64^Cu and Exo-NOTA-^64^Cu.

**SUPPLEMENTAL FIGURE 5.**
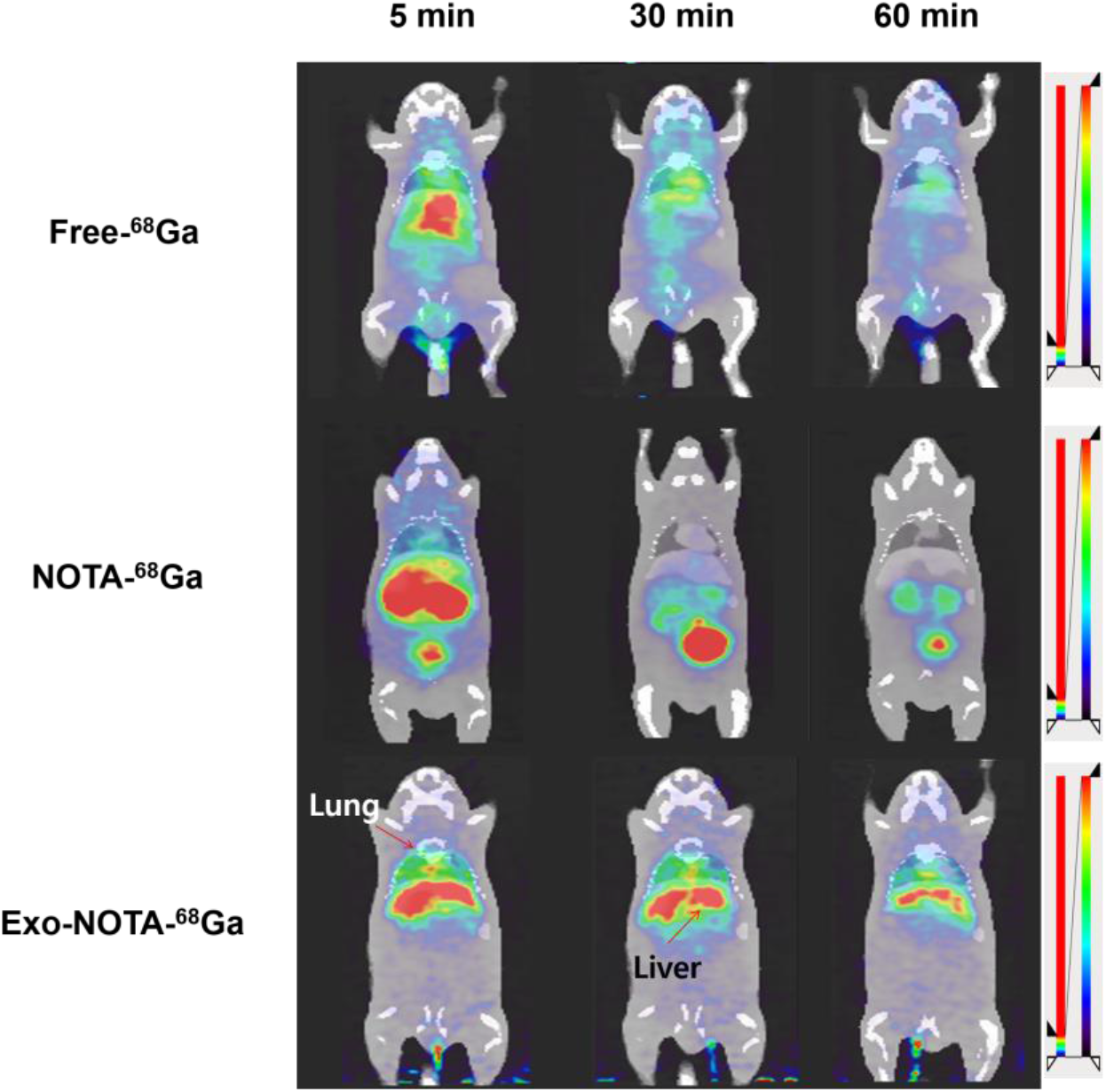
^68^Ga-labeled exosomes in the hematogenous route. Exo-NOTA-^68^Ga is compared with Free-^68^Ga and NOTA-^68^Ga.

